# Functional idiosyncrasy has a shared topography with group-level connectivity alterations in autism

**DOI:** 10.1101/2020.12.18.423291

**Authors:** Oualid Benkarim, Casey Paquola, Bo-yong Park, Seok-Jun Hong, Jessica Royer, Reinder Vos de Wael, Sara Lariviere, Sofie Valk, Danilo Bzdok, Laurent Mottron, Boris Bernhardt

## Abstract

Autism spectrum disorder (ASD) is commonly understood as a network disorder, yet case-control analyses against typically-developing controls (TD) have yielded somewhat inconsistent patterns of results. The current work was centered on a novel approach to profile functional network idiosyncrasy, the inter-individual variability in the association between functional network organization and brain anatomy, and we tested the hypothesis that idiosyncrasy contributes to connectivity alterations in ASD. Studying functional network idiosyncrasy in a multi-centric dataset with 157 ASD and 172 TD, our approach revealed higher idiosyncrasy in ASD in the default mode, somatomotor and attention networks together with reduced idiosyncrasy in the lateral temporal lobe. Idiosyncrasy was found to increase with age in both ASD and TD, and was significantly correlated with symptom severity in the former group. Association analysis with structural and molecular brain features indicated that patterns of functional network idiosyncrasy were not correlated with ASD-related cortical thickness alterations, but closely with the spatial expression patterns of intracortical ASD risk genes. In line with our main hypothesis, we could demonstrate that idiosyncrasy indeed plays a strong role in the manifestation of connectivity alterations that are measurable with conventional case-control designs and may, thus, be a principal driver of inconsistency in the autism connectomics literature. These findings support important interactions between the heterogeneity of individuals with an autism diagnosis and group-level functional signatures, and help to consolidate prior research findings on the highly variable nature of the functional connectome in ASD. Our study promotes idiosyncrasy as a potential individualized diagnostic marker of atypical brain network development.

## INTRODUCTION

Autism spectrum disorder (ASD) is one of the most common and persistent neurodevelopmental conditions. Behaviorally diagnosed on the basis of clinical observations and standardized tools assessing atypical communication, social interaction, and sometimes restricted and repetitive behaviors and interests (APA, 2013), the broad umbrella term of ASD has resulted in a steady increase in autism prevalence (Baio et al., 2018). This increase in diagnostic sensitivity has on the other hand led to an increasing recognition of heterogeneity of diagnosed individuals (Hong et al., 2018; Hong et al., 2020a; Lombardo et al., 2019), and challenges for specificity (Mottron and Bzdok, 2020). This high variability is present at the phenotypic level of behavioral symptoms and at the level of genetic mechanisms previously associated with ASD (Bernard Paulais et al., 2019; Iuculano et al., 2020; Jeste and Geschwind, 2014), and renders the study of autism particularly challenging. As etiology and pathophysiology remain largely unclear and similarly heterogeneous, efforts have increasingly shifted to neuroimaging techniques to identify intermediary autism phenotypes (Bernhardt et al., 2017; Hong et al., 2019). It is hoped that these can potentially consolidate molecular perturbations and behavioral perspectives on ASD and identify biomarkers of symptom severity.

Fuelled by the increased availability of data sharing initiatives (Di Martino et al., 2017; Di Martino et al., 2014; Milham et al., 2018; Poldrack et al., 2017), numerous neuroimaging studies based on resting-state functional magnetic resonance imaging (rs-fMRI) have indicated that autistic individuals often present with a mosaic pattern of connectivity alterations between distributed cortical regions relative to typically developing (TD) controls (Assaf et al., 2010; Di Martino et al., 2014; Keown et al., 2013; Monk et al., 2009; von dem Hagen et al., 2013). These connectivity alterations often manifest in the form of connectivity reductions in both higher order association cortices as well as sensory and motor regions, and sometimes co-occur with patches of connectivity increases between cortical and subcortical nodes (Cerliani et al., 2015). However, other research has also emphasized *i*) little overlap between reported results, *ii*) variable patterns of hyper/hypo-connectivity, and *iii*) an impact of preprocessing choices as well as subject-specific head motion and other confounds on observed findings (Cerliani et al., 2015; Cheng et al., 2017; Holiga et al., 2019; King et al., 2019; Lynch et al., 2013; Müller et al., 2011). Inconsistent findings have also been attributed to the conventional use of case-control designs in connectomics research in autism, which assume within-group homogeneity (Lenroot and Yeung, 2013; Zabihi et al., 2019). In addition to efforts that attempt to address this heterogeneity by subtyping ASD individuals into more homogeneous groups (Hong et al., 2020a; Lombardo et al., 2019), a nascent literature has emphasized the importance to study inter-individual variability of functional connectivity patterns in ASD compared to TD (Dickie et al., 2018; Hahamy et al., 2015; Nunes et al., 2019). This body of work suggests that such idiosyncrasy may be an important feature of functional connectome organization in ASD, with greater variability in functional topography among ASD individuals relative to TD. At the group level, this may potentially impact the analysis of connectivity differences between ASD and TD when assuming an identical alignment between the functional and structural domains among individuals. In other words, anatomical alignment does not guarantee a correspondence of intrinsic functional profiles. Ignoring this phenomenon may lead to losing subject-specific features of network organization at the group level (Braga and Buckner, 2017; Poldrack, 2017). In ASD, given the highly idiosyncratic nature of the functional connectome, this is even more pronounced (Hahamy et al., 2015), leading to spurious differences in connectivity that might be better explained when taking into consideration this heterogeneity (Byrge et al., 2015; Castles et al., 2014).

Although recent work has suggested an idiosyncratic organization of the functional connectome in ASD (Dickie et al., 2018; Hahamy et al., 2015; Nunes et al., 2019), here we expand these approaches in several important ways. First, we developed a novel multi-marker profiling of idiosyncrasy, based on measures of spatial variability, connectome manifold analysis and well as probabilistic approaches to characterize uncertainty of subject-specific functional topographies. These descriptors comprehensively profiled differences in idiosyncrasy between ASD and TD, and provided the bases for an assessment of associations to age and symptom severity. To furthermore identify structural and potential molecular factors that give rise to the spatial patterns of ASD-related network idiosyncrasy, we correlated idiosyncrasy findings in ASD against MRI-based cortical thickness findings as well as *post-mortem* gene expression information. Indeed, prior research has demonstrated atypical cortical development in ASD (Khundrakpam et al., 2017; Zielinski et al., 2012), with genetic risk factors likely to play a major role in brain anatomy and connectivity abnormalities (Abrahams and Geschwind, 2010). Finally, we tested our main hypothesis, and assessed how idiosyncrasy may relate to connectivity alterations in ASD vis-a-vis healthy controls observed at the group level (Bzdok et al., 2020; Bzdok and Meyer-Lindenberg, 2018). Specifically, we conducted a group-level analysis to study functional connectivity differences between ASD and TD, capitalizing on prior graph theoretical measures (Holiga et al., 2019), with and without considering idiosyncrasy.

## RESULTS

We studied idiosyncrasy based on rs-fMRI data from both waves of the Autism Brain Imaging Data Exchange (ABIDE I and II) (Di Martino et al., 2017; Di Martino et al., 2014), a multi-site data-sharing initiative. Specific site inclusion criteria and rigorous data quality control as in prior work (Hong et al., 2019; Park et al., 2020b; Valk et al., 2015) resulted in a total of 329 participants (157/172 ASD/TD) from 5 different sites (see **Tables S1-S2**). Our image processing strategy involved the mapping of functional signals to cortical surfaces as well as surface-based spherical alignment, on which functional connectivity matrices were calculated at a single-subject level. Diffusion map embedding, a non-linear dimensionality reduction technique that projects regions into a low-dimensional space governed by similarity in connectivity profiles (Coifman and Lafon, 2006; Margulies et al., 2016), identified a common low-dimensional manifold where individual embeddings were clustered into seven intrinsic connectivity networks (ICNs) using a Gaussian mixture model. Connectivity idiosyncrasy was characterized with two complementary features, namely the analysis of spatial shifting on the cortical surface meshes and the analysis of dispersion in connectome-based manifolds. Descriptors were computed relative to a reference embedding (and its corresponding clustering) built by averaging all individual connectivity matrices (see **Figure 1A**). Findings corrected for site, age, and sex unless otherwise specified. Further information about the dataset, image processing, and idiosyncrasy descriptors is provided in the *Methods* section.

**Figure 1.**
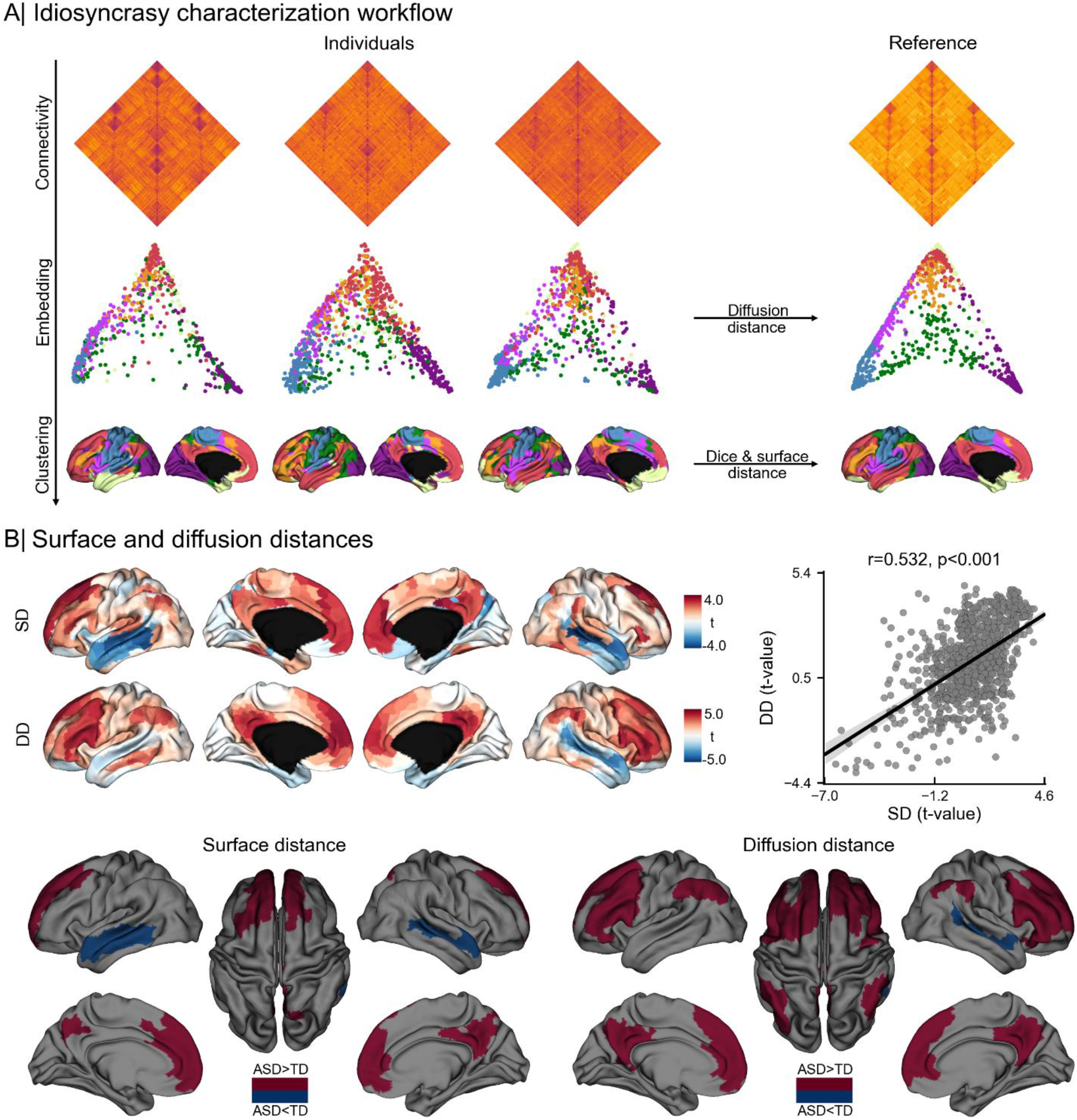
Spatial shifting and diffusion distance of intrinsic connectivity networks in ASD and TD. A) Proposed workflow for intrinsic connectivity identification and idiosyncrasy characterization. B) Statistical t-maps of surface (SD) and diffusion (DD) distance differences and their Pearson’s correlation (top), and areas showing significant idiosyncrasy differences in SD and DD (bottom).

### Idiosyncrasy is characterized by shifting of functional networks in physical and embedding spaces

Idiosyncrasy was assessed through the quantification of surface distance (SD) and diffusion distance (DD). In brief, SD is the geodesic distance from a given point to the closest point in the corresponding reference network (see **Figure 1**A). DD, on the other hand, profiles idiosyncrasy in terms of the similarity in the connectivity patterns across individuals and with the canonical reference in the embedding space (see *Idiosyncrasy descriptors* section). Both approaches showed increased idiosyncrasy in ASD relative to TD in medial and lateral prefrontal regions in both hemispheres, with DD showing more marked effects bilaterally in the precuneus and angular gyrus (see **Figure 1**B). Interestingly, ASD also showed bilateral reductions in idiosyncrasy compared to TD in lateral temporal cortices.

We complemented the surface-based analysis with an assessment of idiosyncrasy at the network-level to determine if observed differences are related to spatial variability or to differences in the size of the ICNs. Here, we computed mean surface distance (MSD) to compare the locations of each ICN in each individual to its corresponding reference network (see **Figure S1**). Higher MSD values indicate that a given individual network deviates from the corresponding reference network. In both ASD and TD, the visual network (VN) showed the least idiosyncratic organization (i.e., lowest MSD), whereas the ventral attention network (VAN) had the most idiosyncratic organization (i.e., highest MSD). These results indicate that idiosyncrasy is network specific. Comparing groups, we found significant differences after FDR correction in the dorsal attention (DAN, p=0.044), default mode (DMN, p=0.005), somatomotor (SMN, p=0.038) and VAN (p=0.001) networks, with ASD showing increased MSD relative to TD. Across the whole cortex, individuals with ASD also showed higher spatial shifting than TD (p=0.004). Similar findings were also obtained when quantifying spatial shifting using Dice overlap measures. As higher MSD (and lower Dice) may also indicate that individual networks span larger/smaller portions of the cortex relative to the reference network because of hyper- or hypo-connectivity, we also analyzed between-group differences in network size. Notably, however, we did not find significant differences suggesting that findings were due to idiosyncrasy rather than connectivity differences per se (see **Table S3**).

We further contextualized idiosyncrasy using a probabilistic framework for each ICNs as shown in **Figure 2**. Qualitatively, we observed more spreading of the probability maps in ASD at the group-level, particularly in DMN and SMN (see **Figure 2**A). This spreading is manifested as higher entropy in ASD (see **Figure 2**B). Increases in ASD were highest in the SMN (Cohen’s *d*=0.290), followed by VAN (*d*=0.209), DAN (*d*=0.203), and DMN (*d*=0.188). On the other hand, the limbic system (LSN, *d*=-0.282) showed lower entropy in ASD (see **Figure 2**C). We could observe high correlations of group-wise entropy differences with the corresponding differences in SD (r=0.512, p_spin_<0.001) and DD (r=0.459, p_spin_<0.001) (see **Figure 2**D), even after accounting for spatial autocorrelation. In accordance with the SD and DD findings, we observed lower entropy in ASD relative to TD in the lateral temporal lobe.

**Figure 2.**
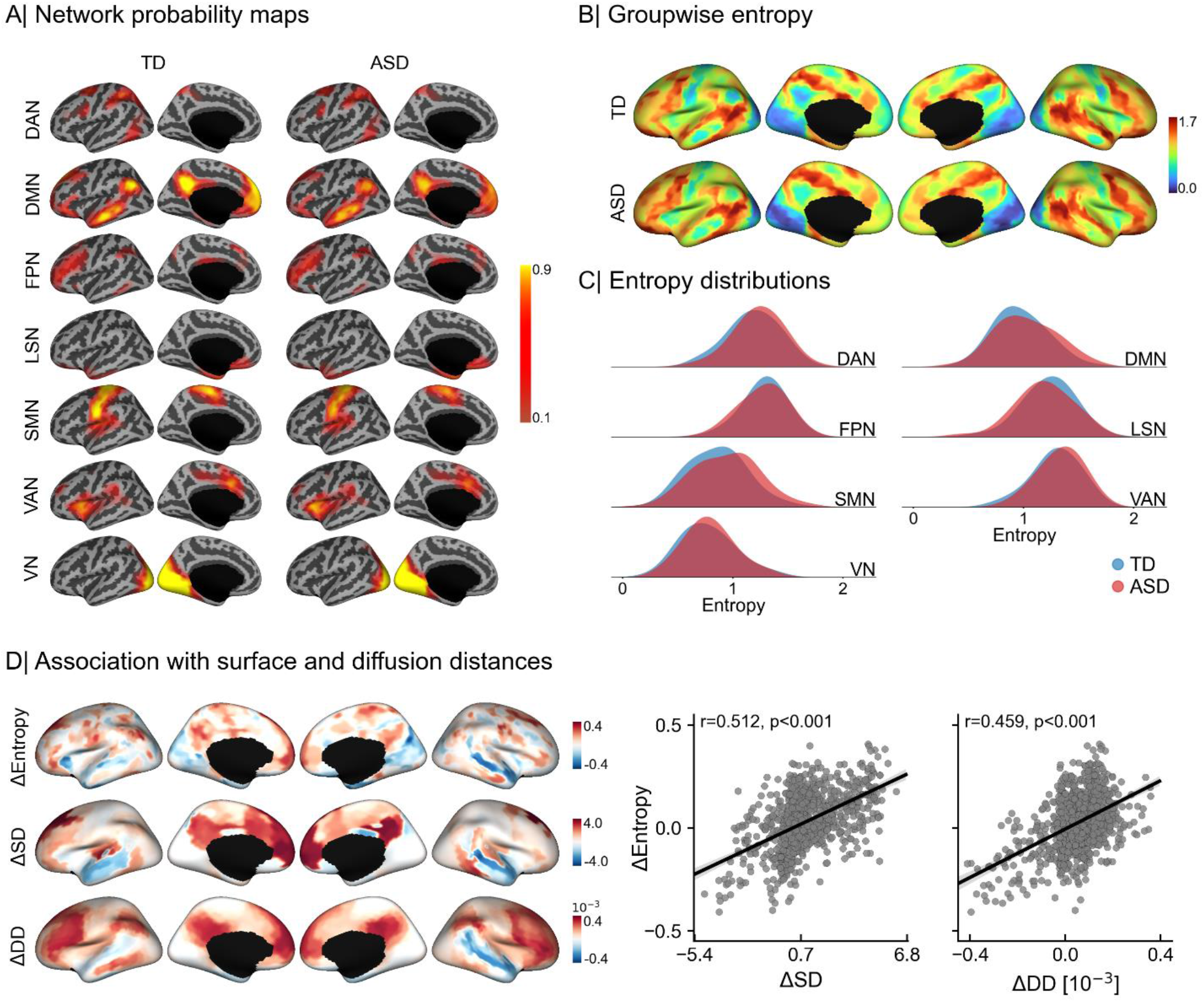
Spatial probability maps and entropy distributions of intrinsic connectivity networks. A) Lateral and medial views of the left hemisphere displaying group-level spatial probability maps for each intrinsic connectivity network in TD (left) and ASD (right). B) Group-wise entropy computed from the average probability maps for TD (top) and ASD (bottom). C) Entropy distributions for each functional network (based on the reference clustering). D) Spatial correlation of group-wise entropy differences (i.e., ΔEntropy) with the corresponding group-wise differences in surface (ΔSD) and diffusion (ΔDD) distances. Positive values reflect higher entropy/distance in ASD.

Although site was included as a covariate in all our analyses, we repeated the SD and DD analyses for each site separately. **Figure S2** and **Figure S3** in supplementary material respectively display global and site-specific SD and DD difference between TD and ASD. Despite variability in findings across sites, the overall direction of findings was relatively consistent across most of the included sites, particularly those with the highest numbers of subjects i.e. NYU, USM, PITT (see **Table S2**).

### Idiosyncrasy association to age and symptom severity

When analyzing age effects (see Figure 3 and **Figure S4**), we found significant associations to idiosyncrasy in the DAN (p<0.001/0.001 for SD/DD), LSN (p=0.089/0.001), SMN (p<0.001 for both SD and DD), VAN (p=0.016/0.001) and VN (p<0.001). On the other hand, we found no significant relationship with DMN (p=0.551/0.423) and FPN (p=0.143/0.087). Overall, there was a significant effect of age on shifting in cortical and embedding spaces (p<0.001), manifested in increasing SD and DD. These results indicate that idiosyncrasy increases with age. Nevertheless, ASD and TD showed similar slopes and there were no significant group-level interactions. We further assessed these results repeating our surface-based analysis using only the children (i.e., age <18 years) in our dataset and only the adults (i.e., age ≥18). Overall, results from these analyses, reported in **Figure S5**, were consistent with the findings obtained when using all individuals in our dataset. Nonetheless, when only using adults, the cluster in the temporal lobe showing higher idiosyncrasy in TD was relatively larger with respect to the cluster found in children.

**Figure 3.**
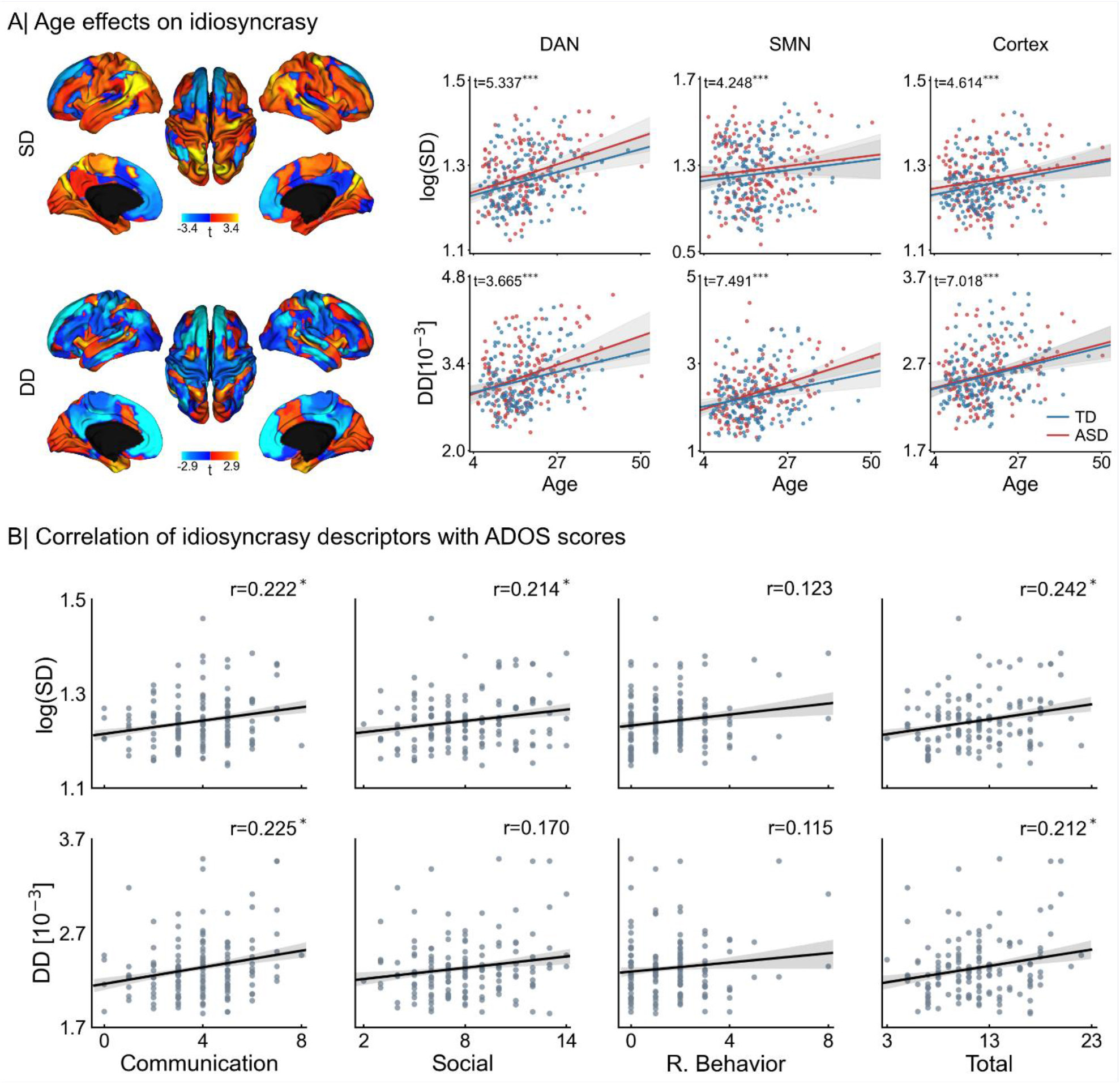
Idiosyncrasy association with age and symptom severity. A) T-maps of age effects on surface and diffusion distances (left), and relationships of surface (top) and diffusion (bottom) distances with age for dorsal attention (DAN) and somatomotor (SMN) networks, and globally for the entire cortex (right). B) Pearson’s correlation of average surface (top) and diffusion (bottom) distances with ADOS scores. Statistical significance is indicated with *, ** and ***, respectively denoting p<0.05, p<0.01, and p<0.001 after FDR correction across 7 networks for age and, for ADOS, across the 4 different scores.

We also investigated the association of our idiosyncrasy descriptors with ASD symptom severity. Specifically, we tested whether Autism Diagnostic Observation Schedule (ADOS) scores were associated with surface and diffusion distances (see **Figure 3**B). The descriptors were computed for the entire cortex. After correcting for multiple comparisons, we found significant associations with communication subscores (SD: r=0.222, p=0.030, and DD: r=0.219, p=0.013), social cognition subscores (SD: r=0.214, p=0.017) and total ADOS scores (SD: r=0.242, p=0.011, and DD: r=0.205, p=0.020). From these results we can see that increasing idiosyncrasy is related to symptom severity.

### Associations to cortical morphology and gene expression patterns

Several studies (Bedford et al., 2020; Hong et al., 2018; Khundrakpam et al., 2017; Pereira et al., 2018; Valk et al., 2015) have reported morphological alterations in ASD relative to TD. Cortical thickness changes were overall consistent with morphological anomalies reported in the literature, showing a mix of frontal and midline parietal cortical thickening together with patches of cortical thinning in temporal regions (Bedford et al., 2020; Khundrakpam et al., 2017; Pereira et al., 2018; Valk et al., 2015; van Rooij et al., 2018). Importantly, we also inspected the relationship of the functional idiosyncrasy descriptors with changes in cortical morphology, by running spatial correlation analyses between group-wise differences in cortical thickness to those in SD and DD. To account for spatial autocorrelations, we used spin tests with 1000 permutations (Alexander-Bloch et al., 2018). As shown in **Figure 4**A, we found no significant associations between cortical thickness and either measure of idiosyncrasy, suggesting that differences in functional idiosyncrasy are not spatially overlapping with potential alterations in cortical morphology in the ASD sample studied here.

**Figure 4.**
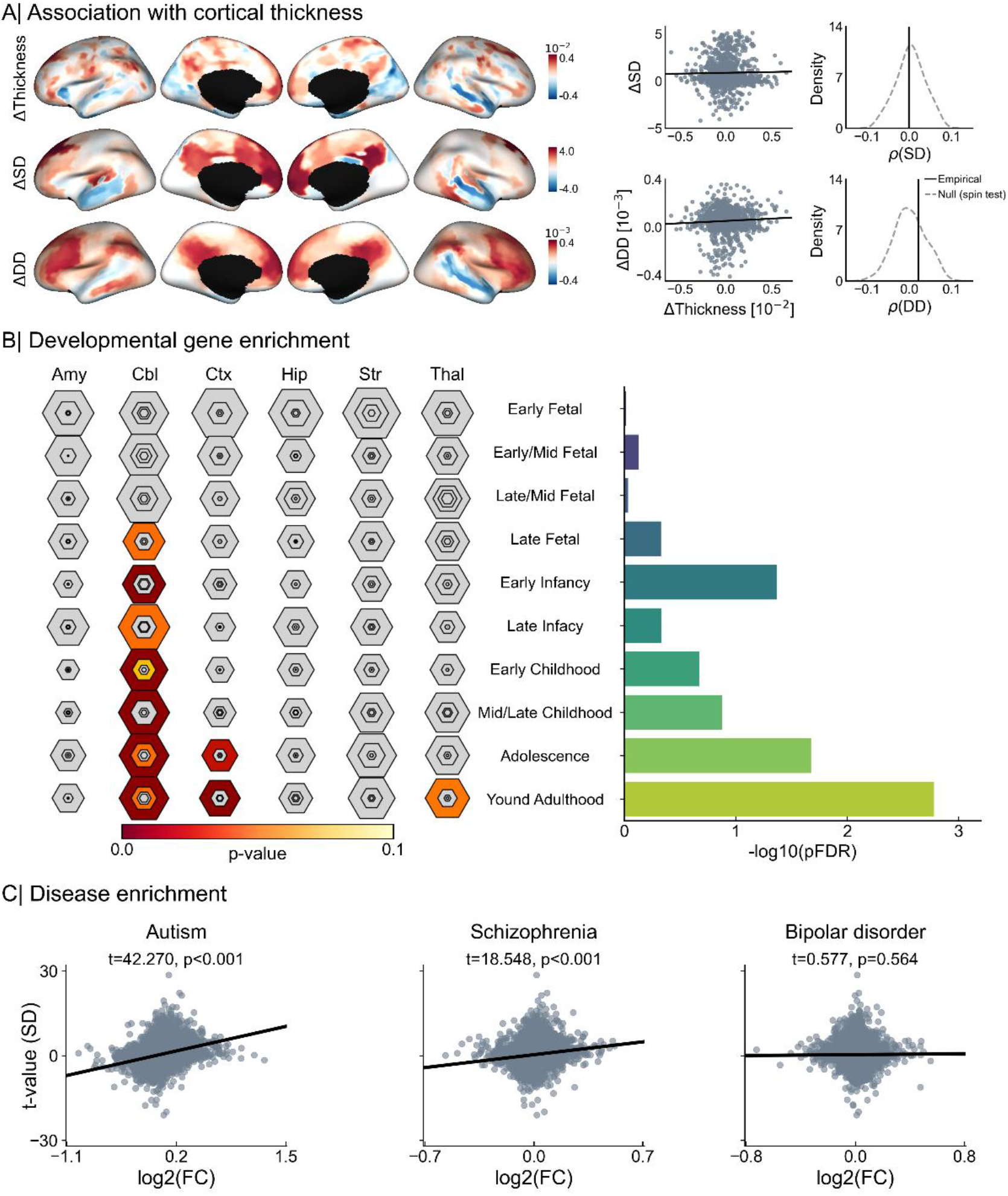
Associations to cortical morphology and gene expression patterns. A) Correlation of group-wise differences in surface (top) and diffusion (bottom) distances with cortical thickness, and comparison of empirical correlation with the null distribution obtained using 1000 spin permutation tests to account for spatial autocorrelation. B) Developmental cortical enrichment, showing enrichment mainly in cerebellum, cortex and striatum (left), specifically in adolescence and young adulthood (right). C) Associations of gene expression in neuropsychiatric disorders, where log2(FC) stands for log2 fold-change of the genes in each disorder and the vertical axis indicates the significance of the relationship strength of the genes with idiosyncrasy in terms of surface distance (SD).

Furthermore, we explored potential neurobiological correlates of idiosyncrasy in ASD. Idiosyncrasy maps obtained using SD and SD were correlated with *post-mortem* gene expression maps from six donors provided by the Allen Institute for Brain Sciences (AIBS) (Hawrylycz et al., 2012), as above. Significant genes were identified by spatially correlating their expression patterns with our maps of idiosyncrasy based on spin tests with 1000 permutations, across the six *post-mortem* cortical brain samples. Only genes that were significantly associated with idiosyncrasy and consistently expressed across all six donors (average inter-donor correlation ≥0.5) were considered for further analysis (Arnatkevič iūté et al., 2019). Selected genes (see **Table S4**) were tested for developmental expression analysis across different developmental time windows (Dougherty et al., 2010), from early fetal to young adulthood, and disease enrichment analysis. This analysis showed associations of our idiosyncrasy descriptors with genes expressed in the brain from early infancy onwards (see **Figure 4**B), and across several brain regions comprising the cerebellum, cortex and striatum. Significant gene expressions were predominantly found in adolescence and young adulthood. Moreover, disease enrichment analysis (see **Figure 4**C and **Figure S6**) revealed that cortical patterns of idiosyncrasy were more strongly associated with differential gene expression (Gandal et al., 2018) in ASD (t=42.270/28.099, p<0.001 for SD/DD) than in schizophrenia (t=18.548/14.192, p<0.001) or bipolar disorder (t=0.577/-2.014, p=0.564/0.044).

### Connectivity alterations are driven by idiosyncrasy

Prior research has suggested connectivity alterations in ASD relative to controls, but patterns of findings have overall not been consistent (Dickie et al., 2018; Hahamy et al., 2015). Here, we examined the relationship between idiosyncrasy (in terms of SD and DD) and overall connectivity alterations, quantified using degree centrality (DC; see below for findings using eigenvector centrality). DC provides an unbiased depiction of the functional connectome that assigns each cortical location the number of connections exceeding a predefined threshold, set here to 0.2 (Holiga et al., 2019).

The relationships of degree centrality with surface and diffusion distances are shown in **Figure 5**A and reported in **Table 1** for each ICN. DC showed strong correlations with both SD (r=0.468, p<0.001) and DD (r=0.413, p<0.001). For DC, positive/negative values indicate hyper/hypo-connectivity in ASD, and higher/lower spatial deviations in surface and diffusion distances relative to TD. These results show that connectivity alterations in ASD are significantly associated with idiosyncrasy. Regions that exhibit hyper-connectivity (i.e., higher DC) in ASD show increased spatial deviation from the locations of the canonical networks. At the network-level, SD was associated with DC in FPN, DMN and VAN, whereas DD was associated with DC in all networks except the limbic system. Given the relationship of idiosyncrasy with DC, we set to investigate the role of idiosyncrasy in the connectivity alterations observed in previous works. To do so, we first analyzed the differences in DC between ASD and TD and then repeated the same analysis controlling for idiosyncrasy. That is, we used both SD and DD as additional covariates in our analysis. As shown in **Figure 5**B, the number of clusters showing significant differences is considerably reduced, with only one small region in the left frontal lobe remaining. Findings were replicated using a different centrality measure (i.e., eigenvector centrality (Bonacich, 2007)), as shown in the supplementary **Figure S7** and **Table S5**. Eigenvector centrality assigns each node its corresponding entry in the eigenvector with the largest eigenvalue of the connectivity matrix. With eigenvector centrality, none of the regions showing significant differences in connectivity survived after controlling for idiosyncrasy. Altogether, these findings suggest that identifiable connectivity alterations in conventional ASD to control comparisons do, at least in part, emanate as a result of the high variability in the spatial locations of the ICNs.

**Figure 5.**
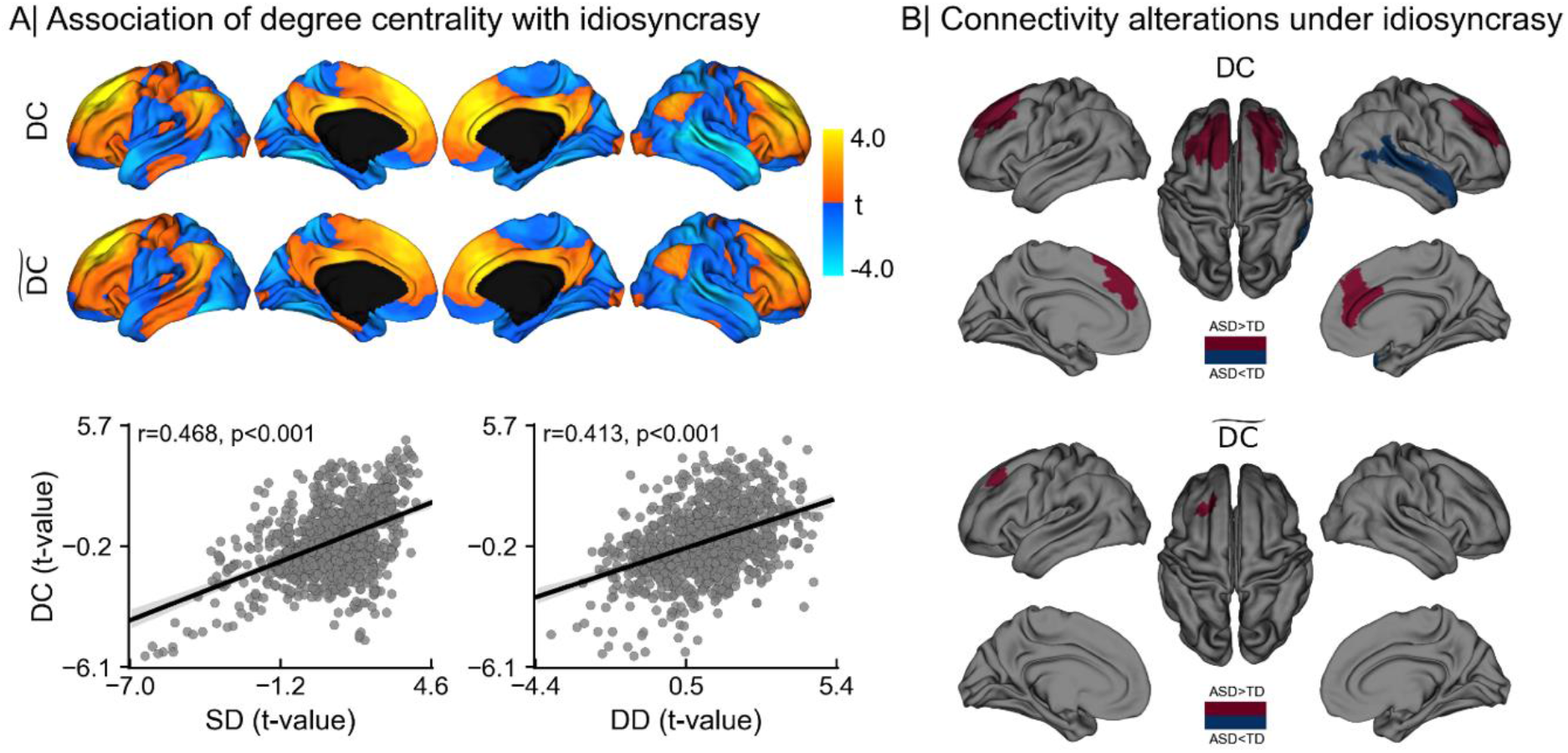
Association of degree centrality with idiosyncrasy. A) Statistical t-maps (top) of areas showing differences in degree centrality between ASD and TD before (i.e., DC) and after controlling for idiosyncrasy (i.e., 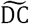), and Pearson’s correlation (bottom) of DC t-map with t-maps of surface (SD) and diffusion (DD) distances. B) Regions showing significant DC increases (red) and decreases (blue) in ASD before (top) and after (bottom) controlling for idiosyncrasy. Idiosyncrasy is represented with SD and DD as additional covariates.

**Table 1.**
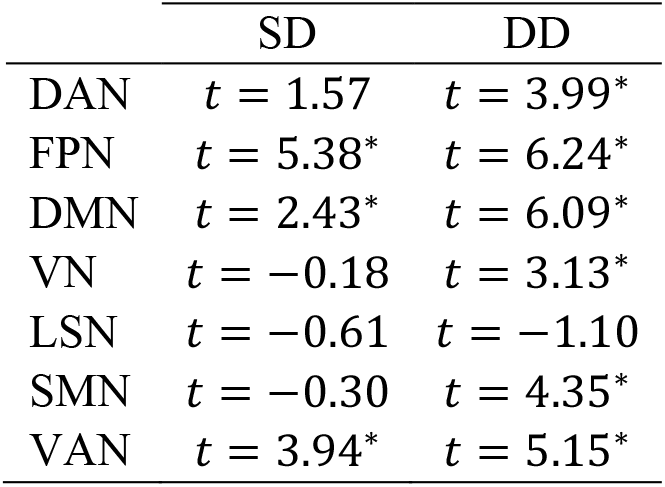
Relationship of idiosyncrasy, in terms of network-wise surface (SD) and diffusion (DD) distances, with average degree centrality for each intrinsic connectivity network. Significant associations after FDR correction are denoted with *.

## DISCUSSION

Neurodevelopment is a complex yet coordinated process shaping the anatomy and function of multiple brain networks, with important variability across individuals. Characterizing this variability may add precision in the study of typical development and may advance our understanding of atypical neurodevelopment in diverse indications such as ASD (Collin and van den Heuvel, 2013; Di Martino et al., 2014). Multiple studies have previously reported atypical functional connectivity in ASD, contributing to the overall notion of ASD as a disorder of brain networks (Di Martino et al., 2014; Holiga et al., 2019; Hong et al., 2019). However, there have also been reports questioning the consistency of findings, both in terms of which networks are involved and in terms of the directionality of findings (He et al., 2020; Müller et al., 2011). Beyond an increasing recognition on the impact of preprocessing choices and sample inclusion criteria (Hong et al., 2020a; Mottron and Bzdok, 2020; Müller et al., 2011), a growing research line is hinting at a more variable and idiosyncratic organization of the functional connectome in ASD as a potential contributor to these inconsistent findings (Dickie et al., 2018; Hahamy et al., 2015; Nunes et al., 2019). In essence, idiosyncrasy describes an increased spatial variability in the mapping between functional network organization and brain anatomy. Here, we set out to *i*) characterize such idiosyncratic network organization in ASD, using novel metrics that capture network variation in both physical and topological spaces, *ii*) examine associations to age and symptom severity, *iii*) explore morphological and genetic associations, and

*iv*) investigate how idiosyncrasy may contribute to functional connectivity alterations commonly seen in ASD to control case-comparison studies. In short, our findings suggest that ASD presents with a mosaic of idiosyncrasy alterations relative to TD, with mainly increases in ASD, together with focal decreases in network idiosyncrasy. Idiosyncrasy was found to relate to both age and symptom load as measured with the ADOS, and the spatial topography of ASD-related network idiosyncrasy strongly correlated with the expression of autism risk genes. Notably, we could also show that idiosyncrasy contributes to ASD versus TD connectivity differences that are detectable with typical case-control analysis, motivating future research strategies that consider patterns of idiosyncrasy in their analyses.

Core to our work were two complementary approaches to quantify functional idiosyncrasy, with one approach operating in the spatial domain and another one in connectivity-determined manifold spaces. Both approaches converged in showing that while functional network organization is idiosyncratic in both TD and ASD, the latter showed a mosaic of mainly increases in network idiosyncrasy across multiple functional systems together with patches of idiosyncrasy reductions. In the spatial domain, we compared individual network locations to a canonical reference connectome, built by averaging all individual connectivity matrices in our dataset. This highlighted that several networks (i.e., DAN, DMN, SMN and VAN) were shifted from the typical locations of their corresponding reference networks in ASD. Complementing idiosyncrasy profiling in the spatial domain, we characterized idiosyncrasy in a connectivity-informed manifold space. Such manifolds provide coordinate systems based on intrinsic network organization and are, thus, decoupled from the underlying anatomy (Hong et al., 2020b; Huntenburg et al., 2018; Margulies et al., 2016). In several recent studies, our group and others capitalized on manifold spaces to represent structural and functional connectome information (Bajada et al., 2020; Paquola et al., 2019; Park et al., 2020a), to assess structure-function coupling (Park et al., 2020a), and to study typical and atypical connectome organization (Burt et al., 2018; Hong et al., 2019; Paquola et al., 2019; Park et al., 2020b; Shine et al., 2019). Unlike their spatial counterparts, manifold-based idiosyncrasy measures tap into inter-subject correlations (Simony et al., 2016), and in turn, provide a metric sensitive to regional connectivity, as well as similarity in connectivity to other regions (Piella, 2014). This descriptor converged with our surface distance analysis, in that it pointed to a spatially varying pattern of idiosyncrasy, with ASD showing mainly increased idiosyncrasy in sensory and higher order networks. In addition to the convergence in findings across these two descriptors, we could cross-validate our findings using an entropy-based descriptor at the network level. This approach provided an independent probabilistic context to understand our findings in terms of inter-individual spatial network uncertainty, supporting increased idiosyncrasy in ASD in multiple networks relative to TD.

The functional networks found to be idiosyncratic in our analyses have been consistently shown to diverge in analyses that compared functional connectivity in ASD relative to TD at the group level. Indeed, several studies have reported connectivity alterations in ASD individuals relative to TD in the DAN (Bi et al., 2018; Farrant and Uddin, 2015; Yerys et al., 2017), VAN (Farrant and Uddin, 2015; Yerys et al., 2019), DMN (Bi et al., 2018; Di Martino et al., 2009; Kozhemiako et al., 2020; Lombardo et al., 2019) and the SMN (Cerliani et al., 2015; Di Martino et al., 2014; Holiga et al., 2019; Kozhemiako et al., 2020). Our findings show that the degree of spatial shifting, irrespective of the cohort, is distributed across the putative functional hierarchy, affecting primary sensory, unimodal association and attentional, as well as higher order transmodal systems such as the DMN. Interestingly, and previously unreported in ASD, our two idiosyncrasy measures also pointed to reduced idiosyncrasy in the lateral temporal lobe in autism. A prior rs-fMRI study in neurotypical individuals (Mueller et al., 2013) found the lateral temporal cortex to be among the areas with the highest inter-subject variability in intrinsic functional connectivity. Moreover, lateral temporal cortical areas have previously been suggested to show abnormal structural connectivity in ASD in very young children at risk for ASD (Lewis et al., 2017). In that study, lower structural network efficiency of primary and secondary auditory cortices was related to autism risk in children as young as 6 months old, and network inefficiencies were related to symptom load at a later follow-up (Lewis et al., 2017). The authors suggested that atypical organization in sensory systems in autism may manifest early, and potentially cascade into the organization of higher order networks – a finding in line with the sensory first hypothesis of autism and other neurodevelopmental disorders (Marco et al., 2011; Mottron et al., 2006; Robertson and Baron-Cohen, 2017).

Findings of increased idiosyncratic organization in ASD are consistent with prior work reporting higher inconsistency in the incorporation of individual anatomical locations to the DMN and SMN (Nunes et al., 2019), and increased spatial shifting in DAN and VAN (Dickie et al., 2018). In our study, all these four ICNs had a more idiosyncratic functional organization in ASD. Moreover, and similar to prior work, we found that idiosyncrasy increased with symptomatology, more specifically in social and communication difficulties, indicating that functional network reorganizations which diverge most from the normative group are reflected in more pronounced patterns of behavioural disturbance on standardized testing. Nonetheless, these prior studies have largely overlooked the relationship of the underlying spatial topography to connectivity differences in ASD versus control populations. In (Hahamy et al., 2015), it was shown that the existence of topographical distortions among individuals leads to a *regression to the mean* effect at the group level. In other words, the study of functional connectivity at the group level may be affected by latent misalignments between the functional organization and the underlying anatomy, potentially giving rise to spurious differences. Indeed, our analysis of functional connectivity alterations showed that idiosyncrasy is a potential confounder. Hyper- and hypo-connected regions found in ASD using degree and eigenvector centrality measures show great overlap with previous findings in multiple large-scale datasets using degree centrality (Holiga et al., 2019). However, after controlling for idiosyncrasy (using both surface and diffusion distances as covariates), differences were considerably reduced. A small patch with increased connectivity survived when using degree centrality, whereas with eigenvector centrality no connectivity differences were found. It is plausible that idiosyncratic reorganization in ASD breaks down the functional correspondence between homologous anatomical regions across individuals assumed in case-control studies, and thus challenges inference as well as the interpretation of previously reported connectivity differences. Inter-subject variability in functional connectivity has been shown to be related to the variability in position of functional regions even in normative populations (Li et al., 2019). This is closely related to an emerging literature on precision neuroimaging in healthy populations, where several studies have also shown specific within-subject features of network organization that do not manifest at the group level due to this effect (Braga and Buckner, 2017; Gordon et al., 2017; Gratton et al., 2018; Poldrack, 2017). In addition to potential spatial uncertainty, other findings have also shown that some of the connectivity alterations found in ASD are partially driven by short-term temporal variability (Falahpour et al., 2016). Taken together, our findings suggest a marked influence of network idiosyncrasy on what is detectable with traditional case-control connectivity analyses. As such, they support the development of novel approaches to analyze connectivity differences at the group level, while also considering subject-specific variability, especially in atypical populations such as ASD.

Correlating idiosyncrasy measures with age indicated age-related increase in both ASD and TD, with however no significant differences in trajectories between groups. As such, our results point to an increased functional network idiosyncrasy in ASD relative to controls already be present at an early age, with neither a considerable aggravation nor normalization throughout childhood development, adolescence and early adulthood. Notably, while out inclusion criteria allowed the study of both children and adults with ASD and TD, our youngest participants were 5 years old. In light of emerging studies suggesting connectivity anomalies in very young children with autism (Lewis et al., 2017), it will therefore be of relevance to assess network idiosyncrasy in small kids and infants, and to also model intra-individual trajectories longitudinally. This will offer a more precise understanding of early mechanisms contributing to idiosyncrasy, alongside with a more direct mapping of intra-individual trajectories in idiosyncratic networks.

Although our findings showing a mosaic pattern of increased and decreased idiosyncrasy warrant further investigation, a plausible explanation for this large-scale functional reorganization in ASD together with increased variability in both spatial and connectome-based network embeddings may relate to compensatory plasticity mechanisms, and its imbalance in autism. By integrating genetic, cognitive, and neuroimaging findings, the so-called trigger threshold target model of autism (Mottron et al., 2014) has postulated that ASD may relate to neurodevelopmental disturbances that trigger compensatory reallocations of neural resources in autism. As a result, intact regions assume functions from nearby impaired areas. To accommodate this shifting of competences, spatially adjacent networks might be required to adjust their locations and/or typical functional crosstalk, a phenomenon that may contribute to the observed increases in SD and DD in the cohort with autism. In this direction, prior work has suggested an abnormal cortical plasticity in ASD (Pedapati et al., 2016), with multiple genetic factors involved in this process. In fact, most genetic risk factors associated with ASD appear to be implicated in synaptic plasticity and connectivity more generally (Bourgeron, 2015; Sestan and State, 2018). These risk factors may be shared across a range of neuropsychiatric disorders (Moreau, 2020). The vast genetic diversity associated with ASD may therefore account for heterogeneity in connectivity alterations observed in ASD (Zerbi et al., 2020). In this work, we investigated the relationship of idiosyncrasy with gene expression, showing that genes associated with idiosyncrasy differences were more strongly correlated with differential gene expression in ASD than in schizophrenia and bipolar disorder, which further highlights idiosyncrasy as an important feature of autism. Note, however, that gene expression used in this analysis is derived from adult *post-mortem* data and our findings may thus only represent indirect associations that need to be confirmed as additional resources and datasets become available.

To conclude, our work characterized functional idiosyncrasy in spatial and connectivity-informed manifold dimensions of the functional connectome. Studying a large dataset of TD and ASD, our novel descriptors reliably captured differences in both groups, suggesting a mosaic pattern of idiosyncrasy increases and decreases in several functional networks in ASD. In addition to showing associations to age, symptom severity, as well as gene expression patterns, our findings notably indicated a marked relationship between idiosyncrasy and connectivity differences that can be identified using case-control analysis, which may consolidate some of the heterogeneity observed in previous studies in ASD and calls for the consideration of idiosyncrasy when studying the functional connectome in autism, since connectivity alterations may, at least partly, reflect an underlying idiosyncratic organization.

## METHODS

### Participants and data acquisition

We studied rs-fMRI data from both waves of the openly-shared Autism Brain Imaging Data Exchange initiative (ABIDE I and II; http://fcon_1000.projects.nitrc.org/indi/abide) (Di Martino et al., 2017; Di Martino et al., 2014). For our study, we selected those sites with ≥10 individuals per group and with both children and adults. After detailed quality control, only cases with acceptable T1-weighted (T1w) MRI, surface-extraction, and head motion in rs-fMRI were included in our analyses, resulting in a total of 329 subjects (157/172 ASD/TD, with mean±SD age in years = 18.4±8.2/18.4±7.7) from 5 different sites: (1) NYU Langone Medical Center (NYU, 35/51 ASD/TD from ABIDE-I, and 21/19 from ABIDE-II); (2) University of Utah, School of Medicine (USM, 49/37 ASD/TD); (3) University of Pittsburgh, School of Medicine (PITT, 19/20 ASD/TD); (4) Trinity Centre for Health Sciences, Trinity College Dublin (TCD, 12/16 ASD/TD); and (5) Institut Pasteur/Robert Debré Hospital (IP, 11/21 ASD/TD). High-resolution T1w images and rs-fMRI were acquired on 3T scanners from Siemens (NYU, USM, PITT) or Philips (IP, TCD). More information about acquisition settings for each site is provided in **Table S1**.

Individuals with ASD were diagnosed by an in-person interview with clinical experts and gold standard diagnostics of the Autism Diagnostic Observation Schedule, ADOS (Lord et al., 2000) and/or Autism Diagnostic Interview-Revised (ADI-R) (Lord et al., 1994). TD individuals did not have any history of mental disorders. For all groups, participants who had genetic disorders associated with autism (i.e., Fragile X), contraindications to MRI scanning, and pregnancy were excluded. The ABIDE data collections were performed in accordance with local Institutional Review Board guidelines, and data were fully anonymized. Detailed demographic information from participants included in our study are reported in **Table S2**.

### Data preprocessing

T1w MRI data were preprocessed with FreeSurfer v5.1 (Dale et al., 1999; Fischl, 2012; Fischl et al., 1999). The pipeline performed automated bias field correction, registration to stereotaxic space, intensity normalization, skull-stripping, and tissue segmentation. White and pial surfaces were reconstructed using triangular surface tessellation and topology-corrected. Surfaces were inflated and spherically registered to *fsaverage*. For the rs-fMRI, we used preprocessed data previously made available by the Preprocessed Connectomes initiative (http://preprocessed-connectomes-project.org/abide). The preprocessing was performed with C-PAC (https://fcp-indi.github.io) and included slice-time correction, head motion correction, skull stripping, and intensity normalization. The rs-fMRI data were de-trended and nuisance effects related to head motion, white matter and cerebrospinal fluid signals were removed using CompCor (Behzadi et al., 2007), followed by band-pass filtering (0.01-0.1 Hz). Finally, rs-fMRI and T1w data were coregistered in MNI152 space using linear and non-linear transformations. Individual rs-fMRI data were mapped to the corresponding mid-thickness surfaces, resampled to the Conte69 template (https://github.com/Washington-University/Pipelines), and smoothed using a 5 mm full-width-at-half-maximum (FWHM) kernel. All segmentations and surfaces were visually inspected. Subjects with erroneous segmentations or framewise displacements greater than 0.3 mm were excluded from our analyses.

### Identification of intrinsic connectivity networks

To identify and quantify idiosyncrasy in functional network organization, we mapped the rs-fMRI data to a low-dimensional space using the following steps (see **Figure 1A**). First, we built the connectivity matrices from the rs-fMRI time-series of each individual in our dataset using linear correlation coefficients. The connectivity matrices were based on a functional parcellation with 1000 labels (Schaefer et al., 2017), Fisher’s z-transformed and thresholded to only keep the 10% of the most similar entries per row (Vos de Wael et al., 2020). We used diffusion mapping introduced in (Coifman and Lafon, 2006), as implemented in BrainSpace (Vos de Wael et al., 2020), to embed the rs-fMRI data into a low-dimensional manifold. This approach is robust to noise and computationally efficient compared to other non-linear manifold learning techniques (Tenenbaum et al., 2000; von Luxburg, 2007). Briefly, diffusion mapping embeds the data into a particular Euclidean space in which the usual Euclidean distance corresponds to the diffusion distance on the data at a given scale or diffusion time. In this new space, interconnected cortical regions are non-linearly projected to fall close to each other, whereas weakly connected regions are mapped to distant locations in the eigenspace. For our study, the diffusion time was set to 1, and the α parameter, which controls the influence of the density of sampling points on the manifold (from maximal influence α=0, to no influence at all α=1) was set to α=0.5 to retain the global relations between data points in the embedded space, following prior work (Hong et al., 2019; Margulies et al., 2016; Vos de Wael et al., 2018). Since diffusion maps capture the main structures of the data along a few cardinal dimensions, we selected the first 30 eigenvectors similar to a previous study, corresponding to the largest eigenvalues to represent each individual embedding (Langs et al., 2016).

To assess differences between TD and ASD, we averaged the connectivity matrices of all the individuals in our dataset to build a mean connectivity matrix, which was subsequently used to construct a reference embedding. This reference embedding was used as a representation of the canonical functional connectivity template. Because diffusion mapping may take the individual datasets into different Euclidean spaces, the standard Euclidean distance between the elements of these spaces is not meaningful. To bring the data into the same Euclidean space, we used a change of basis operator to map all the individual embeddings to the reference embedding (Coifman and Hirn, 2014). In this way, we can compute the Euclidean distance within and between datasets, allowing us therefore to compare the individual diffusion maps to the reference embedding.

Finally, to identify the ICNs, all embeddings (including the reference) were clustered into 7 components using a Gaussian mixture model with full covariance matrix. Each point in the embedding was assigned to the cluster corresponding to the highest a posteriori probability. The mixture model was initialized with the 7 ICNs proposed in (Yeo et al., 2011).

### Idiosyncrasy descriptors

Two different approaches were proposed to characterize idiosyncrasy, namely: spatial- and manifold-based distance measures. For the spatial measure, we used surface distance (i.e., SD), which was computed for each point as the geodesic distance to the closest point in the corresponding reference network (Ecker, et al., 2013; Margulies, et al., 2016; Hong, et al., 2018). The second measure to characterize idiosyncrasy is based on diffusion distance, which is approximated using the Euclidean distance in the eigenspace between points of each individual to the reference embedding, such that points that fall far apart from their corresponding reference points show a high difference in their original rs-fMRI time-series. Instead of computing the distance between pairs of points, however, we take advantage of the clustering and compute the diffusion distance from a given point to its closest point in the reference embedding that belong to the same cluster (i.e., ICN). Let 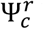 be the set of points of the reference embedding in cluster *c*, for each point 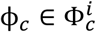 of the individual embedding *i* in the same cluster, our diffusion-based idiosyncrasy descriptor is then computed as follows:

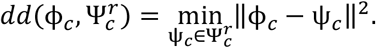

In this way, this descriptor captures the variability that exists in the connectivity patterns that characterize a specific ICN among all the individuals in our dataset. Furthermore, to assess that idiosyncrasy differences are not related to the size of the ICNs, which would rather indicate alterations in connectivity, we aimed to quantify the spatial variability at the network level by computing the overlap and the extent of shifting of each individual clustering from the canonical reference for each ICN. To do so, we used the Dice similarity coefficient (Dice, 1945), which is in common use in neuroimaging research (Eickhoff et al., 2015):

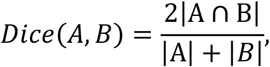

and mean surface distance:

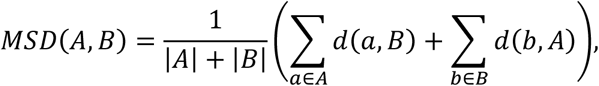

where *A* and *B* are respectively the reference and individual clusters corresponding to a specific network, |·| denotes cardinality, and *d*(*a, B*) is the geodesic distance between point *a* in cluster *A* to the closest point in cluster *B*. In this case, the idiosyncrasy of the individual functional connectomes is indicated by low Dice overlaps and high MSD from the reference clustering.

The spatial variability existing in the network locations among individuals is reflected in the spreading of the ICN probability maps at the group level. The higher the spreading, the more idiosyncratic are the individuals in a given cohort. Therefore, we further characterized idiosyncrasy using a measure of uncertainty based on the entropy of the group-wise probability maps obtained from clustering. In other words, this descriptor measures how evenly the probability mass is spread among the different ICNs at each location. Entropy is minimized when most of the probability mass is concentrated on a particular network, indicating a location with very low variability among individuals. On the other hand, entropy is increased when the probability mass is spread among several ICNs.

### Analysis of idiosyncrasy

Idiosyncrasy was quantified using surface and diffusion distance measures. Group comparisons and correlational analyses additionally controlled for site, sex and age effects. For analyses involving spatial idiosyncrasy descriptors (i.e., Dice overlap and surface distances), surface area was further included as a nuisance covariate. For all our surface-based analyses, threshold-free cluster enhancement (TFCE) was used with 10000 permutations to correct for multiple comparisons across the cortical surfaces (Smith and Nichols, 2009). A significance level of 0.05 was used for all statistical tests.

- Assessing idiosyncrasy differences between ASD and TD. For the spatial descriptors of idiosyncrasy (i.e., Dice overlap and mean surface distance), general linear models (GLM) predicting each of the idiosyncrasy measures based on group diagnosis were used to assess differences at the network level and cortex-wise. For the latter, overall Dice and mean surface distance were computed as the weighted average of the corresponding scores for each ICN, using the size of the reference networks as weights. All results from our network-level analyses were corrected for multiple comparisons using Benjamini-Hochberg FDR correction (Benjamini and Hochberg, 1995). GLMs were also used in the surface-based analysis to study the differences in diffusion and surface distances. For entropy, network-wise differences were analyzed at the group level using two-sample t-tests.
- Age effects. To investigate age-specific differences in idiosyncrasy, the surface-based analysis to study differences in diffusion and surface distances was further repeated separately for children (86 TD and 88 ASD individuals with age <18), and adults (86 TD and 69 ASD individuals with age ≥18) separately.
- Association of idiosyncrasy with ASD symptomatology. Idiosyncrasy descriptors were correlated with ADOS scores, and findings were corrected for multiple comparisons using the FDR procedure. For the correlations of idiosyncrasy with ADOS, we z-scored the data with respect to TD and regressed out the effects of age, sex and site prior to performing the correlations.
- Associations to morphology. To assess whether there is an association between these morphological alterations in ASD and functional idiosyncrasy, we correlated group-wise differences in cortical thickness with surface-based idiosyncrasy measures (i.e., using surface and diffusion distances). We accounted for spatial autocorrelations using non-parametric permutation tests (i.e., spin tests) (Alexander-Bloch, et al., 2018).

### Gene enrichment analysis

Many risk factors have been associated with neurodevelopmental disorders, with genetic factors playing an important role in the etiology of ASD (Miles, 2011; Rylaarsdam and Guemez-Gamboa, 2019). We therefore aimed to investigate the genetic correlates of idiosyncrasy in ASD. Using a similar approach to Neurovault gene decoding tool (Gorgolewski et al., 2015; Hawrylycz et al., 2012), coherent associations between our idiosyncrasy maps (i.e., t-maps of surface and diffusion distances) and *post-mortem* gene expression patterns from the Allen Institute for Brain Sciences (AIBS) were measured to identify the set of genes with significant spatial overlaps. Significant genes were obtained by regressing each gene against our cortical map of idiosyncrasy (e.g., diffusion distance) for each donor and using a one-sample t-test to determine whether the slopes across all six donors were different from 0. To correct for multiple comparisons, the procedure was repeated by randomly rotating our maps of idiosyncrasy using 1000 spin permutations (Alexander-Bloch et al., 2018), which were compared with the original t-statistic to assess gene significance.

Gene expressions for all six donors in the AIBS dataset was obtained using abagen (https://github.com/rmarkello/abagen). Only genes that were consistently expressed across donors (i.e., average inter-donor correlation ≥ 0.5) were considered for our analyses (Arnatkevičiūté et al., 2019). The final set of significant genes was compared against developmental expression profiles from the BrainSpan dataset (http://www.brainspan.org) using the Cell-type Specific Expression Analysis (CSEA) developmental expression tool (http://genetics.wustl.edu/jdlab/csea-tool-2/). Disease enrichment analysis was also conducted based on a recently published catalog of genes associated with neuropsychiatric disorders that share similar genetic variants (Gandal et al., 2018). We used robust linear regression to assess the relationship between the genes using the t-statistics derived from the previous spatial analysis with their fold change of expression in ASD, schizophrenia and bipolar disorder (Gandal et al., 2016). Guanine-cytosine content was used as an additional covariate to control for possible effects related to genome size in microarray data (Love et al., 2016; Steijger et al., 2013).

### Relation to degree centrality

Given the little consensus on the directionality of the connectivity alterations in ASD reported in the literature. Here, our purpose is to investigate the relationship between idiosyncrasy and connectivity alterations to elucidate the role of idiosyncrasy in these connectivity alterations. To study this putative association, we used two different measures of centrality, namely: degree and eigenvector centrality. The first measure is defined as the total number of connections whose linear product moment correlation coefficients are above a predefined threshold used to eliminate connections with low temporal correlation attributable to signal noise (Buckner et al., 2009; Holiga et al., 2019). Eigenvector centrality is based on the eigenvector with the largest eigenvalue of the connectivity matrix. Following prior work that used these measures to study connectivity alterations in ASD (Di Martino et al., 2013; Holiga et al., 2019), the threshold for our analyses was set to 0.2.

Since idiosyncrasy is an inherent property that is also present in TD individuals (presumably in a lower degree than in ASD), we first analyzed the relationship of idiosyncrasy with hyper- and hypo-connectivity based on linear product moment correlations of the statistical t-maps of degree and eigenvector centrality with those of diffusion and surface distances using spin tests (Alexander-Bloch et al., 2018). Positive degree centrality values would indicate hyper-connectivity in ASD, whereas negative values indicate hypo-connectivity. The same applies to our idiosyncrasy descriptors, with positive/negative surface distances, for instance, pointing out to higher/lower deviations from the canonical reference networks relative to TD. Then, we investigated the impact of idiosyncrasy in the potential connectivity alterations when ignoring this phenomenon. Surface-based analysis to find differences in connectivity between ASD and TD was performed based on degree centrality (or eigenvector centrality). This analysis was initially conducted without considering idiosyncrasy and then repeated controlling for idiosyncrasy by incorporating SD and DD as additional covariates to our GLMs.

## ACKNOWLEDGEMENTS

Oualid Benkarim was funded by a Healthy Brains for Healthy Lives (HBHL) postdoctoral fellowship and is a member of the Quebec Autism Research Training (QART) program. Casey Paquola was funded through a postdoctoral fellowship of the Fonds de la Recherche due Quebec - Santé (FRQ-S). Bo-yong Park was funded by the National Research Foundation of Korea (NRF-2020R1A6A3A03037088), Molson Neuro-Engineering fellowship by Montreal Neurological Institute and Hospital (MNI) and FRQ-S. Boris Bernhardt acknowledges research support from the National Science and Engineering Research Council of Canada (NSERC Discovery-1304413), the Canadian Institutes of Health Research (CIHR FDN-154298), SickKids Foundation (NI17-039), Azrieli Center for Autism Research (ACAR-TACC), BrainCanada (Azrieli Future Leaders), and the Tier-2 Canada Research Chairs program. Jessica Royer was funded by a CIHR fellowship. Reinder Vos de Wael was funded by a studentship from the Savoy Foundation. Sara Lariviere is funded by CIHR. We would also like to acknowledge support from the Helmholtz Foundation and the Healthy Brains for Healthy Lives initiative. Danilo Bzdok was supported by the Healthy Brains Healthy Lives initiative (Canada First Research Excellence fund), Google (Research/Teaching Award), the Canadian Institute of Health Research (CIHR), and by the CIFAR Artificial Intelligence Chairs program (Canada Institute for Advanced Research), as well as by NIH-R01 grant AG068563A. Sofie Valk was supported by the Max Planck Society (Otto Hahn Award).

## AUTHOR CONTRIBUTIONS

O.B. and B.C.B. designed the experiments, analyzed the data, and wrote the manuscript. C.P., B.P. and S.H. aided with the experiments. J.R., R.V., S.L., S.V., D.B. and L.M. reviewed the manuscript. O.B. and B.C.B. are the corresponding authors of this work and have responsibility for the integrity of the data analysis.

## COMPETING INTERESTS

The authors declare no conflicts of interest.

## Notes

### Competing Interest Statement

The authors have declared no competing interest.

