## Supplementary material for "Functional idiosyncrasy has a shared topography with group-level connectivity alterations in autism"

*[1]McConnell Brain Imaging Centre, Montreal Neurological Institute and Hospital, McGill University, Montreal, Quebec, Canada; [2]Center for the Developing Brain, Child Mind Institute, New York, NY, USA; [3]Center for Neuroscience Imaging Research, Institute for Basic Science, Sungkyunkwan University, Suwon, South Korea; [4]Department of Biomedical Engineering, Sungkyunkwan University, Suwon, South Korea; [5]Max Planck Institute for Human Cognitive and Brain Sciences, Leipzig, Germany; [6]INM-7, FZ Jülich, Jülich, Germany; [7]Department of Biomedical Engineering, Faculty of Medicine, McGill University, Montreal, Canada; [8]Mila – Quebec Artificial Intelligence Institute, Montreal, Canada; [9]Centre de recherche du CIUSSSNIM et Département de Psychiatrie, Université de Montréal, Montreal, Quebec, Canada*

| | Scanner | Modality | Sequence | TR (ms) | TE (ms) | TI (ms) | FA | matrix | #Vol | Voxel ($mm^3$) |
|---|---|---|---|---|---|---|---|---|---|---|
| IP | Philips | T1w | 3D-MPRAGE | 2500 | 5.60 | | 30° | 240×240 | | 1.0×1.0×1.0 |
| | Achieva | rs-fMRI | 2D-EPI | 2700 | 45.00 | | 90° | 63×63 | 85 | 3.59×3.65×4.0 |
| NYU | Siemens | T1w | 3D-TurboFLASH | 2530 | 3.25 | 1100 | 7° | 256×256 | | 1.0×1.0×1.3 |
| | Allegra | rs-fMRI | 2D-EPI | 2000 | 15.00 | | 90° | 80×80 | 180 | 3.0×3.0×4.0 |
| PITT | Siemens | T1w | 3D-MPRAGE | 2100 | 3.93 | 1000 | 7° | 269×269 | | 1.1×1.1×1.1 |
| | Allegra | rs-fMRI | 2D-EPI | 1500 | 35.00 | | 70° | 64×64 | 200 | 3.1×3.1×4.0 |
| TCD | Philips | T1w | 3D-MPRAGE | 3000 | 3.90 | 1150 | 8° | 256×256 | | 0.9×0.9×0.9 |
| | Achieva | rs-fMRI | 2D-EPI | 2000 | 27.00 | | 90° | 80×80 | 210 | 3.0×3.0×3.2 |
| USM | Siemens | T1w | 3D-MPRAGE | 2300 | 2.91 | 900 | 9° | 240×256 | | 1.0×1.0×1.2 |
| | TrioTim | rs-fMRI | 2D-EPI | 2000 | 28.00 | | 90° | 64×64 | 240 | 3.4×3.4×3.0 |

**Table S1**. Scanner and data acquisition settings. Abbreviations: TR, repetition time; TE, echo time; TI, inversion time; FA, flip angle; #Vol, number of volumes; IP, Institut Pasteur/Robert Debré Hospital; NYU, New York University Langone Medical Center; PITT, University of Pittsburgh, School of Medicine; TCD, Trinity Centre for Health Sciences, Trinity College Dublin; USM, University of Utah, School of Medicine.

| | N (ASD/TD) | Sex (M/F) | Age | ADOS Comm | ADOS Behav | ADOS Social | ADOS Total |
|---|---|---|---|---|---|---|---|
| IP | 11/21 | 18/14 | 20.5±8.8 | 5.3±2.1 | 1.6±2.1 | 9.5±3.5 | 14.8±5.3 |
| NYU | 56/70 | 121/5 | 15.0±7.4 | 3.2±1.7 | 1.8±1.2 | 7.6±2.6 | 10.8±3.9 |
| PITT | 20/22 | 42/- | 20.2±7.1 | 4.2±1.1 | 2.6±1.2 | 8.4±2.3 | 12.7±3.0 |
| TCD | 18/19 | 37/- | 15.2±3.3 | 2.9±0.9 | 0.2±0.5 | 5.8±2.4 | 8.7±2.4 |
| USM | 52/40 | 92/- | 22.7±7.7 | 4.7±1.4 | 1.8±1.8 | 8.9±2.5 | 13.6±3.3 |
| All | 157/172 | 310/19 | 18.4±8.0 | 3.9±1.7 | 1.7±1.5 | 8.1±2.8 | 12.0±4.0 |

**Table S2**. Demographics for each site. First column (N) indicates the number of individuals with autism (ASD) and typically developing (TD); M/F is the number of males/females in each site. Age and Autism Diagnostic Observation Schedule (ADOS) are expressed in mean and standard deviation. Abbreviations: IP, Institut Pasteur/Robert Debré Hospital; NYU, New York University Langone Medical Center; PITT, University of Pittsburgh, School of Medicine; TCD, Trinity Centre for Health Sciences, Trinity College Dublin; USM, University of Utah, School of Medicine. For ADOS: Comm, ADOS communication; Behav, ADOS repeated behavior/interest; and social and total ADOS.

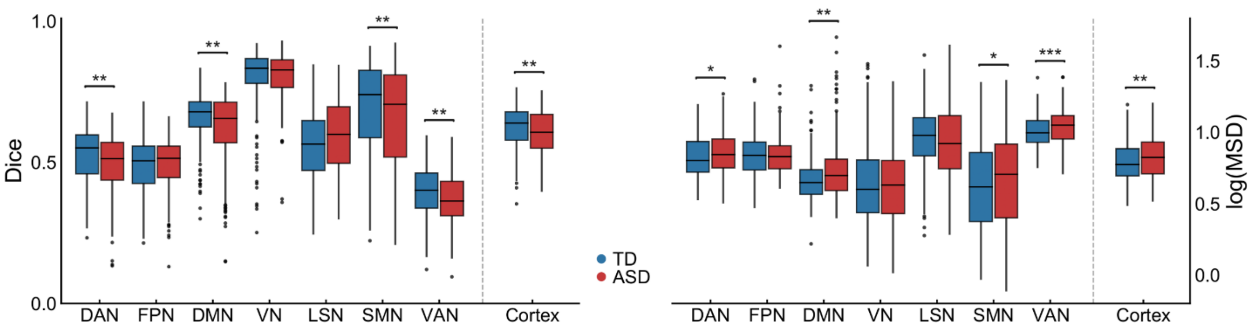


**Figure S1**. Group-wise boxplots of Dice overlap (left) and mean surface distance (MSD) (right) for each intrinsic connectivity network (i.e., cluster) and across the entire cortex. Statistical significance is indicated with *, ** and ***, respectively denoting p<0.05, p<0.01, and p<0.001 after FDR correction across 7 networks.

| | **ICN size** |
|---|---|
| DAN | t=-0.20, p=0.839 |
| FPN | t=2.44, p=0.052 |
| DMN | t=-2.06, p=0.092 |
| VN | t=0.49, p=0.725 |
| LSN | t=2.62, p=0.053 |
| SMN | t=-1.72, p=0.151 |
| VAN | t = 0.64, p = 0.725 |

**Table S3**. Intrinsic connectivity network (ICN) size differences between ASD and TD after controlling for age, sex and site. No statistically significant differences were found.

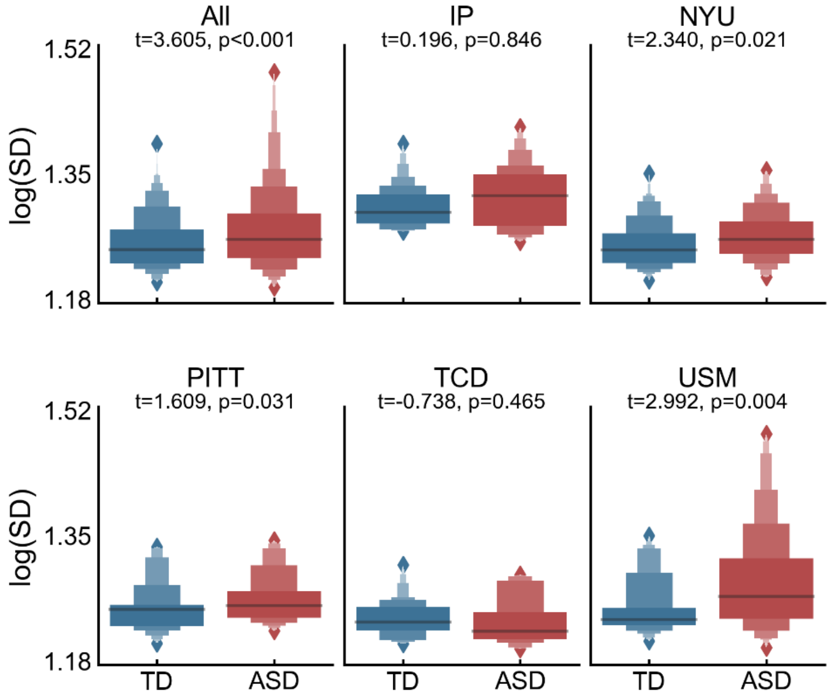


**Figure S2**. Site-specific idiosyncrasy differences. Distributions of mean surface distance in TD and ASD for all sites in the ABIDE I and II datasets and for each site individually.

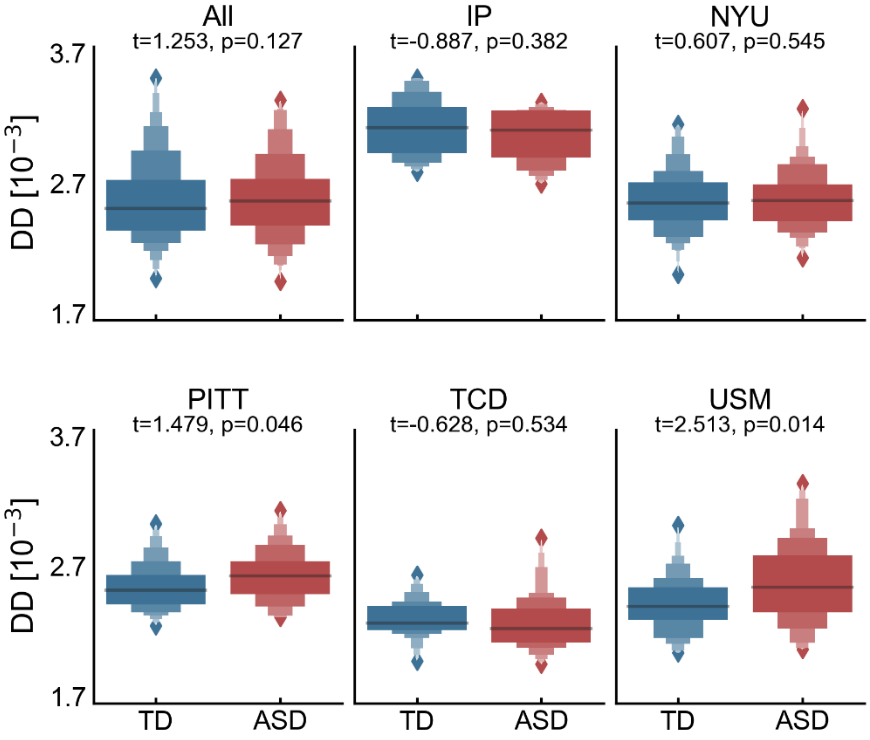


**Figure S3**. Site-specific idiosyncrasy differences. Distributions of mean diffusion distance in TD and ASD for all sites in the ABIDE I and II datasets and for each site individually.

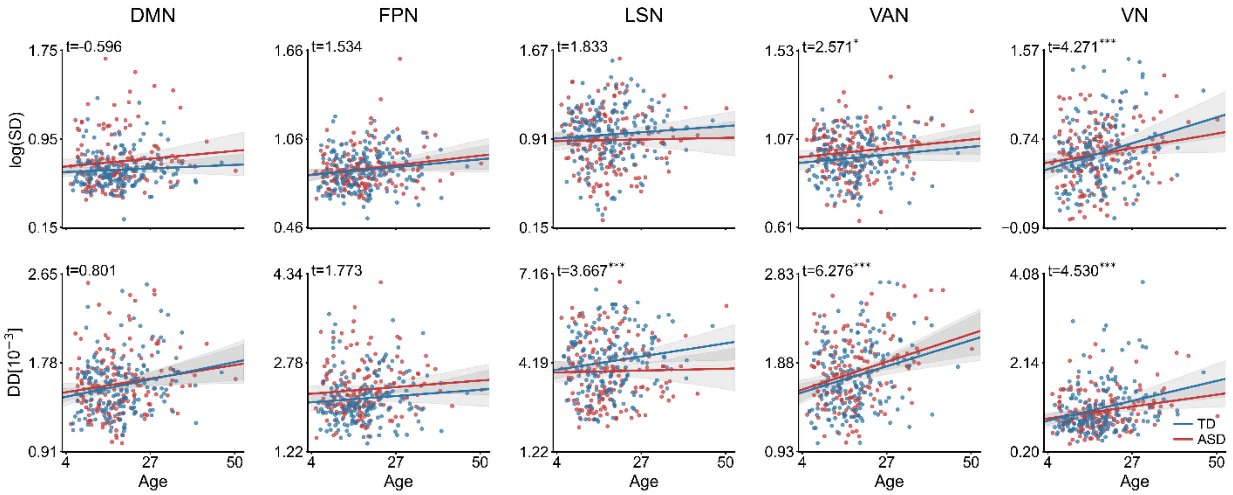


**Figure S4**. Relationship of spatial shifting with age. Average surface (top) and diffusion (bottom) distances in ASD and TD versus age for default mode (DMN), frontoparietal (FPN), limbic system (LSN), ventral attention (VAN) and visual (VN) networks. Remaining networks are available in the main document. Statistical significance is indicated with *, ** and ***, respectively denoting p<0.05, p<0.01, and p<0.001 after FDR correction across 7 networks.

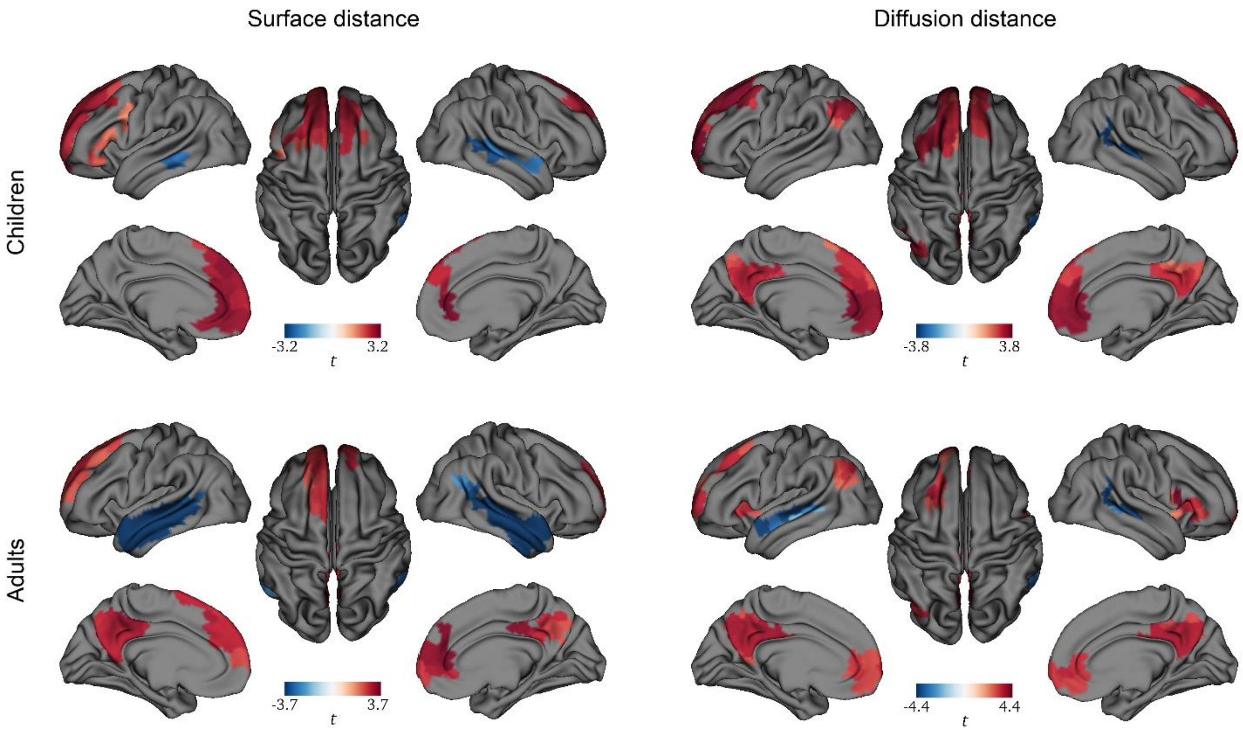


**Figure S5**. Idiosyncrasy differences, using surface (left) and diffusion (right) distances, between ASD and TD in children (i.e., <18 years) and adults (≥18 years). Significance maps obtained using permutation-based thresholding with 10000 permutations.

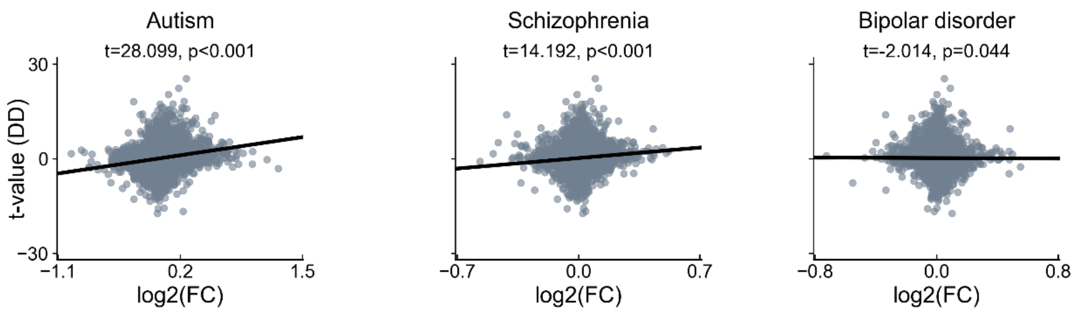


**Figure S6**. Associations of gene expression in neuropsychiatric disorders, where log2(FC) stands for log2 fold-change of the genes in each disorder and the vertical axis indicates the significance of the relationship strength of the genes with idiosyncrasy in terms of diffusion distance (DD).

| **Gene** | **Name** |
|---|---|
| AGO1 | argonaute RISC component 1 |
| ALDH1A3 | aldehyde dehydrogenase 1 family member A3 |
| ANK1 | ankyrin 1 |
| ANKH | ANKH inorganic pyrophosphate transport regulator |
| BEND6 | BEN domain containing 6 |
| CCNO | cyclin O |
| COL5A1 | collagen type V alpha 1 chain |
| CPLX1 | complexin 1 |
| DMKN | Dermokine |
| ECM1 | extracellular matrix protein 1 |
| ESRRG | estrogen related receptor gamma |
| GLCCI1 | glucocorticoid induced 1 |
| GPCPD1 | glycerophosphocholine phosphodiesterase 1 |
| HS3ST1 | heparan sulfate-glucosamine 3-sulfotransferase 1 |
| KCNAB3 | potassium voltage-gated channel subfamily A regulatory beta subunit 3 |
| KCNC1 | potassium voltage-gated channel subfamily C member 1 |
| KCNC3 | potassium voltage-gated channel subfamily C member 3 |
| LINC00599 | MIR124-1 host gene |
| LYSMD4 | LysM domain containing 4 |
| MAGI2-AS3 | MAGI2 antisense RNA 3 |
| MET | MET proto-oncogene, receptor tyrosine kinase |
| MIR124-2HG | MIR124-2 host gene |
| MIR29B2CHG | MIR29B2 and MIR29C host gene |
| NFIC | nuclear factor I C |
| NIPAL2 | NIPA like domain containing 2 |
| NOXO1 | NADPH oxidase organizer 1 |
| OIP5-AS1 | OIP5 antisense RNA 1 |
| RAD54B | RAD54 homolog B |
| SCN1B | sodium voltage-gated channel beta subunit 1 |
| SCRT1 | scratch family transcriptional repressor 1 |
| SEMA7A | semaphorin 7A (John Milton Hagen blood group) |
| SHD | Src homology 2 domain containing transforming protein D |
| SLC25A37 | solute carrier family 25 member 37 |
| UPP1 | uridine phosphorylase 1 |
| WDR97 | WD repeat domain 97 |

**Table S4**. List of genes (symbol and name) significantly associated with idiosyncrasy maps of surface and diffusion distances.

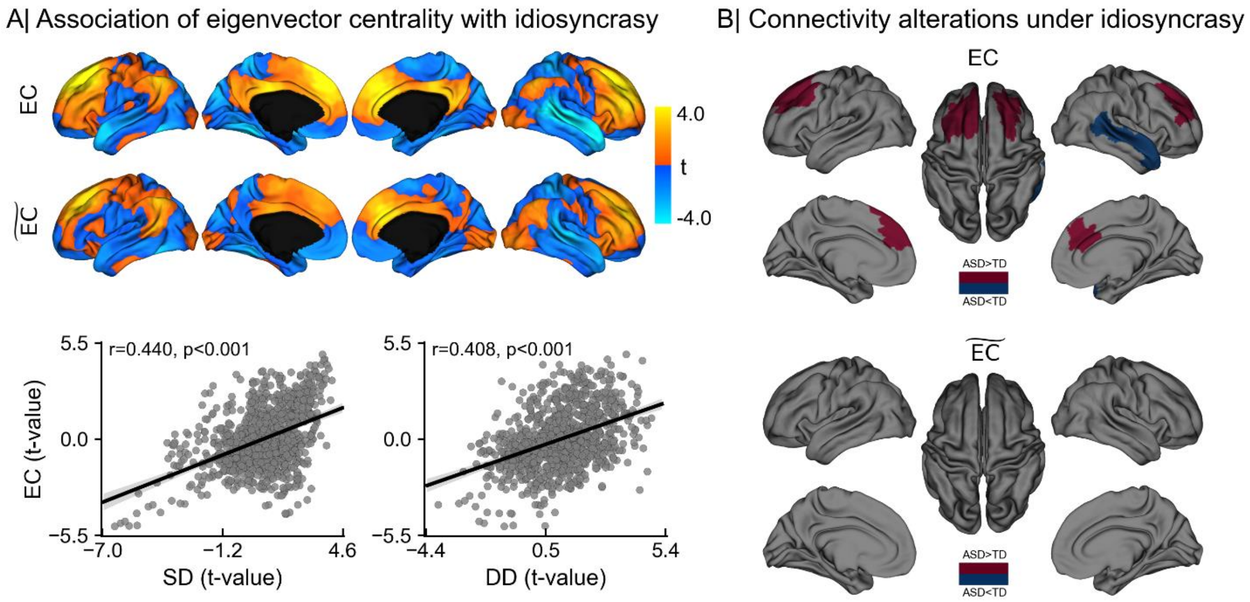


**Figure S7**. Association of eigenvector centrality with idiosyncrasy. A) Statistical t-maps (top) of areas showing differences in eigenvector centrality between ASD and TD before (i.e., EC) and after controlling for idiosyncrasy (i.e., $\widetilde{\mathrm{EC}}$), and Pearson's correlation (bottom) of EC t-map with t-maps of surface (SD) and diffusion distances (DD). B) Regions showing significant EC increases (red) and decreases (blue) in ASD before (top) and after (bottom) controlling for idiosyncrasy. Idiosyncrasy is represented with SD and DD as additional covariates. No significant differences were found after controlling for idiosyncrasy.

| | SD | DD |
|---|---|---|
| DAN | $t = 1.99^*$ | $t = 3.98^*$ |
| FPN | $t = 1.95$ | $t = 5.84^*$ |
| DMN | $t = 3.34^*$ | $t = 5.39^*$ |
| VN | $t = 2.09^*$ | $t = 5.75^*$ |
| LSN | $t = 1.51$ | $t = 4.90^*$ |
| SMN | $t = -2.16^*$ | $t = 4.80^*$ |
| VAN | $t = 1.71^*$ | $t = 3.45^*$ |

**Table S5**. Relationship of idiosyncrasy, in terms of network-wise surface (SD) and diffusion (DD) distances, with average eigenvector centrality for each intrinsic connectivity network. Significant associations after FDR correction are denoted with *.
